# Near infrared spectroscopy detection of hemispheric cerebral ischemia following middle cerebral artery occlusion in rats

**DOI:** 10.1101/2022.08.30.505941

**Authors:** Ardy Wong, Mohammad Iqbal Hossain Bhuiyan, Jeffrey Rothman, Kelly Drew, Kambiz Pourrezaei, Dandan Sun, Zeinab Barati

**Author notes:** **Corresponding author:** Zeinab Barati, PhD, Address: P.O. Box 750198, Fairbanks, AK 99775.

## Abstract

Timely and sensitive in vivo estimation of ischemic stroke-induced brain infarction are necessary to guide diagnosis and evaluation of treatments’ efficacy. The gold standard for estimation of the cerebral infarction volume is magnetic resonance imaging (MRI), which is expensive and not readily accessible. Measuring regional cerebral blood flow (rCBF) with Laser Doppler flowmetry (LDF) is the status quo for confirming reduced blood flow in experimental ischemic stroke models. However, rCBF reduction following cerebral artery occlusion often does not correlate with subsequent infarct volume. In the present study, we employed the continuous-wave near infrared spectroscopy (NIRS) technique to monitor cerebral oxygenation during 90 min of the intraluminal middle cerebral artery occlusion (MCAO) in Sprague-Dawley rats (n=8, male). The NIRS device consisted of a controller module and an optical sensor with two LED light sources and two photodiodes making up two parallel channels for monitoring left and right cerebral hemispheres. Optical intensity measurements were converted to deoxyhemoglobin (Hb) and oxyhemoglobin (HbO_2_) changes relative to a 2-min window prior to MCAO. Area under the curve (auc) for Hb and HbO_2_ was calculated for the 90-min occlusion period for each hemisphere (ipsilateral and contralateral). To obtain a measure of total ischemia, auc of the contralateral side was subtracted from the ipsilateral side resulting in ΔHb and ΔHbO_2_ parameters. Infarct volume (IV) was calculated by triphenyl tetrazolium chloride (TTC) staining at 24h reperfusion. Results showed a significant negative correlation (r = -0.81, p = 0.03) between ΔHb and infarct volume. In conclusion, our results show feasibility of using a noninvasive optical imaging instrument, namely NIRS, in monitoring cerebral ischemia in a rodent stroke model. This cost-effective, non-invasive technique may improve the rigor of experimental models of ischemic stroke by enabling in vivo longitudinal assessment of cerebral oxygenation and ischemic injury.

## 1. Introduction

Stroke is the fifth most common cause of death and a leading cause of long-term disability among adults in the United States. Despite a concerted effort from pharmaceutical companies to develop novel therapies, more than 300 neuroprotective and thrombolytic drug candidates for ischemic stroke have failed in clinical trials between 1995 and 2015 (Chen & Wang, 2016), raising concerns regarding the efficacy of preclinical studies. Several national and international initiatives have been established, including the Stroke Preclinical Assessment Network (SPAN) (Lyden et al., 2022), to address the significant need to improve the scientific investigation of stroke. In addition, the recent successful development of thrombectomy for acute ischemic stroke has generated considerable enthusiasm to reassess neuroprotectant candidates in combination with thrombectomy.

Challenges in translational stroke research is attributable to inconsistent animal models, reliance on short-term end points, and inadequate optimization of therapeutic time window (Fisher et al., 2009; Savitz & Fisher, 2007; Sena et al., 2010; Sutherland et al., 2012). In most animal stroke studies, infarct size is the main outcome measure and any treatment effect is evaluated by statistical comparison of infarct sizes of treatment versus placebo groups. Although parameters of experimental stroke models such as location and duration of vessel occlusion can be standardized, variability in infarct size is yet substantial (standard deviations of ∼30%) due to variability of the surgical procedures and inter-animal variability of blood vessels anatomy (Leithner et al., 2015). Thus, in vivo knowledge of infarct volume before initiating a treatment is important for unbiased exclusion of outliers and reducing variability. It also prevents overestimation (Button et al., 2013) (which is abound in neuroscience publications) or underestimation of treatments’ effects. Diffusion-weighted and perfusion-weighted magnetic resonance imaging (MRI) are sensitive and specific tools for diagnosing infarction after stroke in humans (Neumann-Haefelin et al., 1999; Sorensen et al., 1996). However, MRI equipment is not generally accessible for preclinical studies, especially in smaller research institutes. Also, MRI of animals should be performed under anesthesia, which can itself be neuroprotective, producing biased outcomes (Bleilevens et al., 2013; Kitano et al., 2007); particularly, isoflurane is known to increase cerebral blood flow (CBF) and reduce the infarct size. Laser Doppler flowmetry (LDF) is the most commonly used technique for CBF measurements during the procedures of middle cerebral artery occlusion (MCAO) to ensure a proper and sustained reduction of CBF (Liu et al., 2009), but it cannot predict the subsequent infarct volume.

Ultimately, there is an unmet need for an affordable and easy-to-use device for accurate validation of animal models of ischemic stroke. Such a tool will enable in vivo estimation of ischemic injury and the longitudinal evaluation of treatment efficacy in preclinical models of stroke. In this study, we used a miniaturized optical imaging tool based on Near Infrared Spectroscopy (NIRS) technique (Jöbsis, 1977) for quantitative in vivo estimation of ischemia-induced brain infarction. If successful, this novel technique will support translational efforts to objectively screen and select highly promising treatment candidates for further study in human clinical trials.

## 2. Materials and Methods

### 2.1. Animal preparation

All animal experiments were approved by the University of Pittsburgh Institutional Animal Care and Use Committee and performed in accordance with the National Institutes of Health Guide for the Care and Use of Laboratory Animals. The manuscript adheres to the ARRIVE guidelines for reporting animal experiments.

Male Sprague-Dawley (CD) rats of 9-10 weeks of age were purchased from Charles River Laboratories (Raleigh, NC) and maintained in the University Laboratory Animal Facility until experimental use. All rats were housed in group (2 per case) in a controlled temperature and humidity environment. They were maintained on a 12-h light/12-h dark cycle and provided with food and water *ad libitum*. At the time of experimental use, rats weighed 290-360 g.

### 2.2. Middle cerebral artery occlusion model

Focal cerebral ischemia was induced by transient occlusion of the left middle cerebral artery (MCA) for 90 min as described previously (Bhuiyan et al., 2017). Briefly, under an operating microscope, the left common carotid artery was exposed through a midline incision. Two branches of the external carotid artery (ECA), occipital and superior thyroid arteries, were isolated and coagulated. The ECA was dissected further distally and ligated. The internal carotid artery (ICA) was isolated and carefully separated from the adjacent vagus nerve. The extra-cranial branch of the ICA, the pterygopalatine artery, was then dissected and temporarily ligated. A 22 mm length of silicon-coated nylon filament (size 4-0, native diameter 0.19 mm; diameter with coating 0.41+/- 0.02 mm; coating length 4-5 mm; Doccol Corporation, Sharon, MA) was introduced into the ECA lumen through a puncture. The silk suture around the ECA stump was tightened around the intraluminal nylon suture to prevent bleeding. The nylon suture was then gently advanced from the ECA to the ICA lumen until mild resistance was felt. For reperfusion, the suture was withdrawn 90 min after MCAO to restore blood flow (reperfusion). Body temperature was maintained for the duration of the experiment between 36.5 °C-37 °C with a heating blanket.

### 2.3. Neurological deficit scoring

A neurological deficit grading system (Bhuiyan et al., 2017) was used to evaluate neurological deficit in at 1, and 24 h after reperfusion. The scores are: 0, no observable deficit; 1, forelimb flexion; 2, forelimb flexion and decreased resistance to lateral push; 3, forelimb flexion, decreased resistance to lateral push, and unilateral circling; 4, forelimb flexion and partial or complete lack of ambulation.

### 2.4. Calculation of infarct volume

After 24 hours of reperfusion, rats were anesthetized with 4% isoflurane vaporized in N_2_O and O_2_ (3:2) and decapitated. Brains were removed and two-millimeter coronal slices were made with a rodent brain matrix. The sections were stained for 20 min at 37°C with 1% 2, 3, 5-triphenyltetrazolium chloride monohydrate. Infarct volume and hemispheric swelling were measured using ImageJ software (NIH, Bethesda, MD, USA). Infarct volume was corrected for swelling as described by Swanson et al. (Swanson et al., 1990). Briefly, the sections were scanned, and the ischemic area in each section was calculated by subtracting the non-infarct area in the ipsilateral hemisphere from the total area of the contralateral hemisphere. The infarct areas were summed across all slices and multiplied by the slice thickness (2 mm) giving the total infarct volume (mm^3^).

### 2.5. NIRS Principles and Instrumentation

In the near infrared range (700–900 nm), water, the main ingredient of tissues in vivo, has the lowest light absorption, whereas deoxy-hemoglobin (Hb) and oxy-hemoglobin (HbO2) chromophores are the main absorbers with distinctive absorption characteristics. By choosing two wavelengths in the near infrared spectrum and measuring the attenuation change at two different time points, the relative change in the concentration of Hb and HbO2 molecules can be calculated using the modified Beer-Lambert law (Cope et al., 1988):

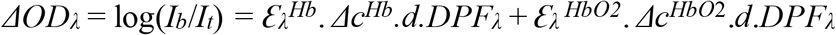

where:

- ΔOD_l_ is optical density, which is the change in optical intensity for the wavelength l;
- I_b_ is the light intensity measured during baseline;
- I_t_ is the light intensity detected during or after a given task;
- 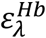 and 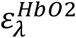 are the absorption coefficients of Hb and HbO2 molecules at the wavelength l;
- Δc^Hb^ and Δc^HbO2^ are the concentration changes of Hb and HbO2 molecules due to the task;
- d is the physical distance between the light source and the photodetector;
- and DPF_l_ is the differential pathlength factor adjusted for the increased pathlength between the light source and the photodetector due to scattering at the wavelength l.

When measured at two wavelengths l_1_ and l_2_, this equation can be solved for the change in the concentration of Hb and HbO2 molecules.

The continuous-wave NIRS system used in this study had three main components: (1) optical sensors consisting of two multiwavelength (730 and 850nm) LED (Marubeni) and two photodiodes (PD15-22C/TR8) built on a flexible circuit substrate to conform to the rat’s head (Figure 1); (2) an Arduino-based controller unit for operating the optical sensors at a 200Hz sampling frequency; (3) a computer running the data collection GUI created in LabVIEW for data acquisition and real-time plotting of the data. The system was calibrated using a solid phantom in accordance with pre-determined specifications.

**Figure 1.**
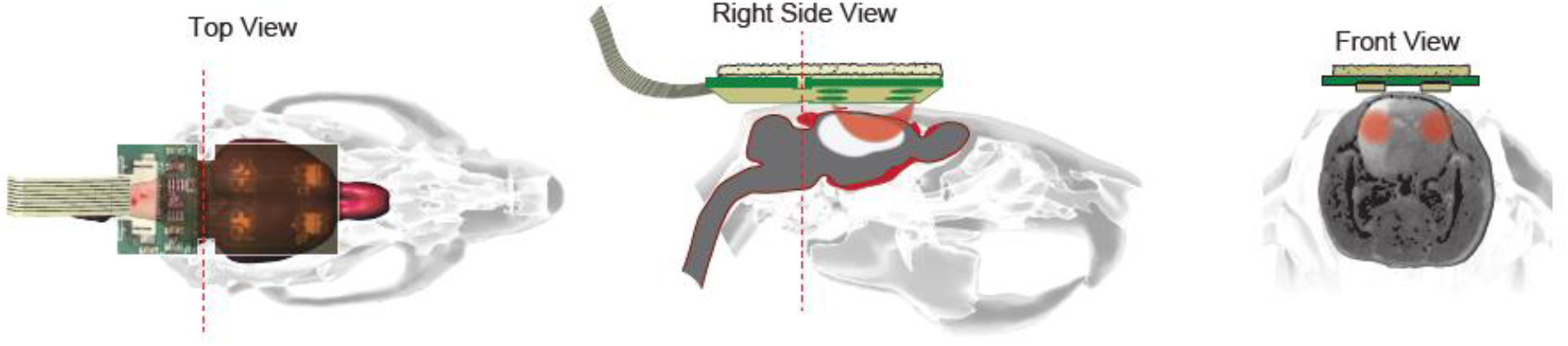
Schematic illustration of the NIRS. Location of the NIRS probe on the rat’s head (Top view). The photons that are emitted by the LED are capturedby the photodiode through the path (orange color banana-shaped path in Right side view). NIRS data is collected on the left and right side of the rat brain to analyze the ipsilateral and the contralateral hemisphere cerebral blood flow (Front view).

### 2.6. Signal processing and data analysis

Signal processing was performed in MATLAB and data analysis was conducted in IBM SPSS. Each channel was visually inspected to reject any saturated or low signals. The accepted NIRS raw data was then filtered through a lowpass filter (fc = 0.1 Hz) where external interferences were present. Area under the curve for the 90-min occlusion period for each hemisphere (ipsilateral and contralateral) were calculated by the trapz function in MATLAB.

## 3. Results

To obtain a measure of total ischemia during 90min occlusion, we subtracted the area under the curve of the contralateral (nonischemic) sides from the ipsilateral (ischemic) side for Hb (ΔHb) and HbO2 (ΔHbO2). Pearson correlation analysis between ΔHb and the infarct volume yielded a significant negative correlation (r = -0.81, p = 0.03); but no significant correlation was found between ΔHbO2 and the infarct volume (r = 0.17, p = 0.70) (Figure 3). In other words, the more deoxygenated the contralateral side relative to the ipsilateral side, the larger the final infarct volume.

The experimental design showing different time points of MCAO, NIRS, neurological scoring and TTC staining is depicted in Figure 2A and representative traces of Hb and HbO2 for two rats with small and large infarct sizes are shown in Figure 2B-C. A neurological deficit score of 3 was obtained in all the seven rats at 1h post-reperfusion. At 24h post-reperfusion, variable scores were obtained with worse scores for animals with larger infarct volumes (Figure 4).

**Figure 2.**
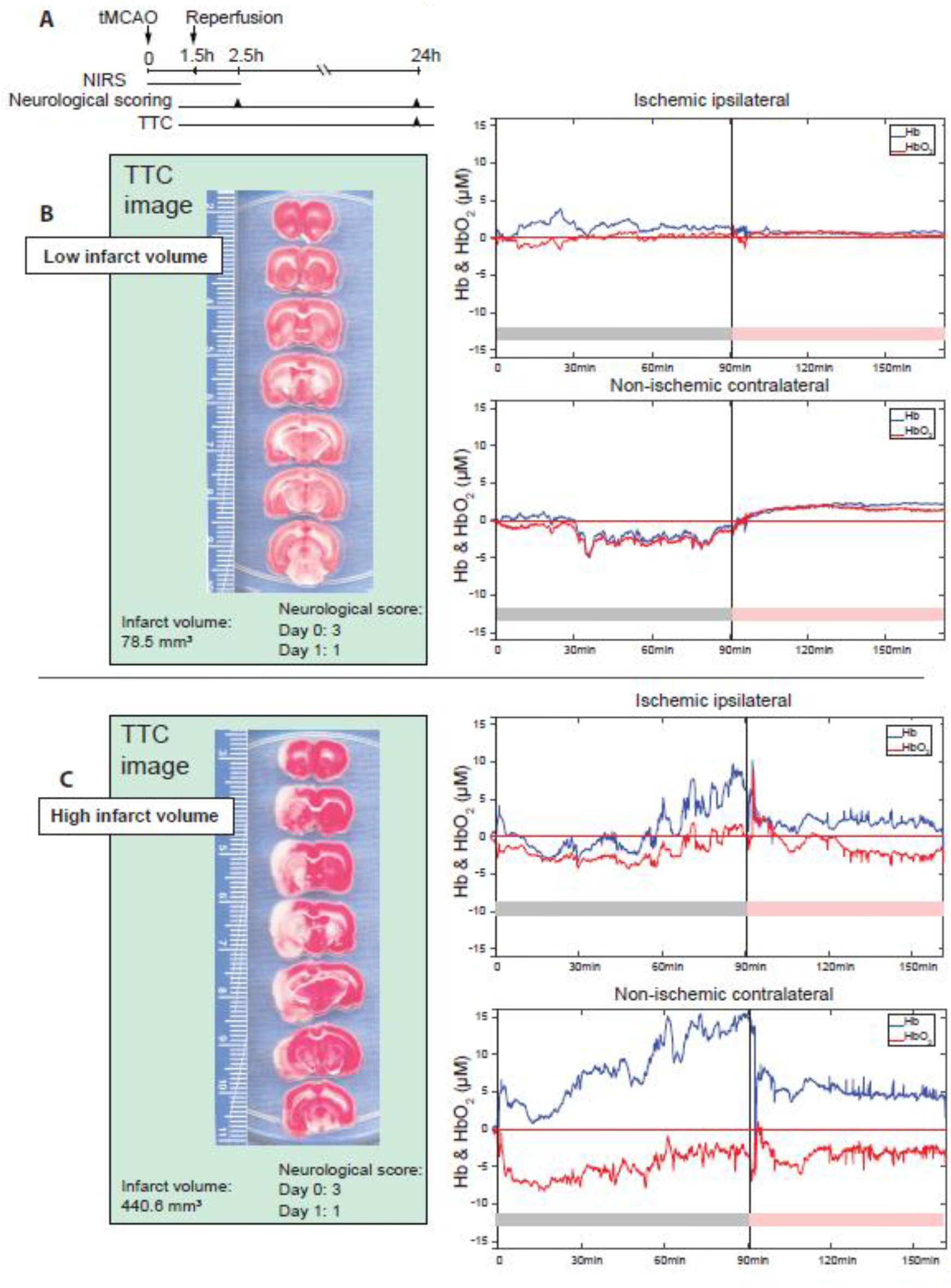
Differences in hemodynamic response during and after stroke. (A) Experimental design showing onset time points of MCAO and NIRS, neurological scoring and TTC staining data collection. (B and C) Representative tracing of continuous NIRS measurement in contralateral (CL) and ipsilateral (IL) hemisphere of rat brain during 90 min of MCAO (grey bar) and 60 min of reperfusion (pink bar). A corresponding image of TTC staining of the same rat’s brain at 24 h after reperfusion is shown. Hb: deoxyhemoglobin, HbO_2_: oxyhemoglobin.

**Figure 3.**
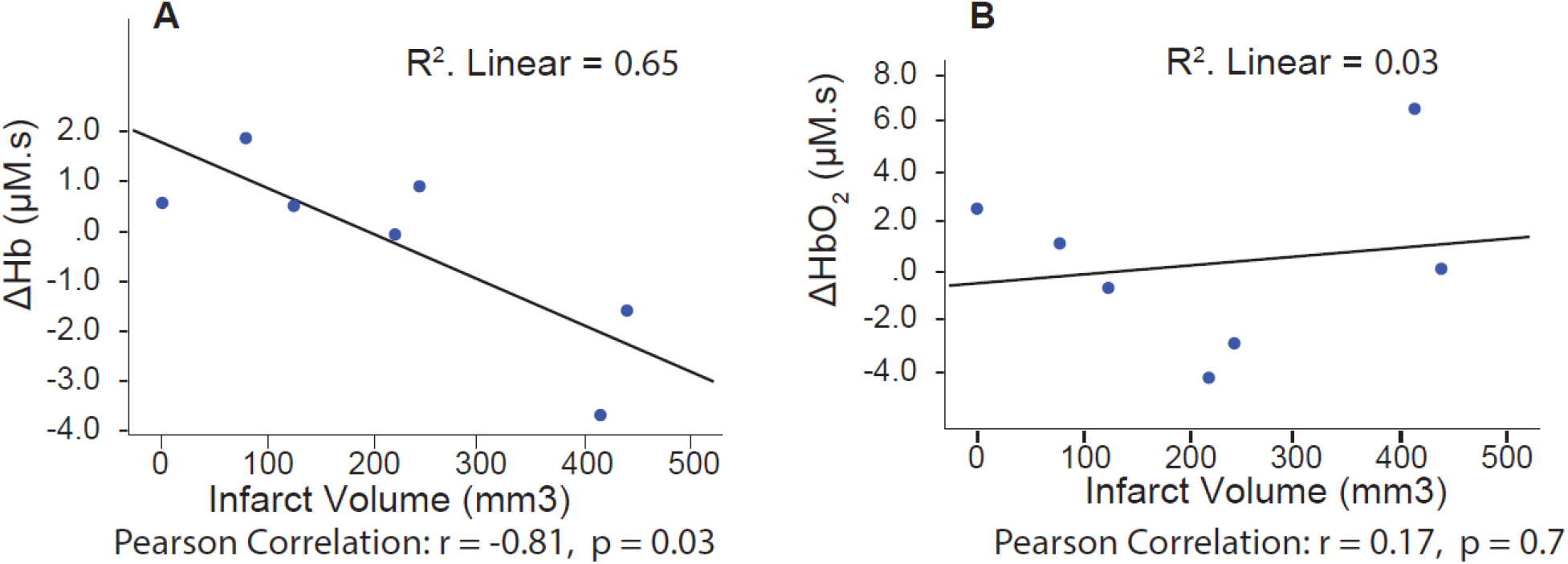
A significant linear correlation was found between ΔHb and infarct volume. Shown are results of correlation analysis between ΔHb, ΔHbO_2_ and infarct volume. Area under the curve (auc) for Hb and HbO_2_ in the 90-min occlusion period from Fig. 2 was calculated for each hemisphere (ipsilateral and contralateral) to obtain a measure of total ischemia for ΔHb (A) and ΔHbO_2_ (B) defined as auc on the contralateral side minus auc on the ipsilateral side.

**Figure 4.**
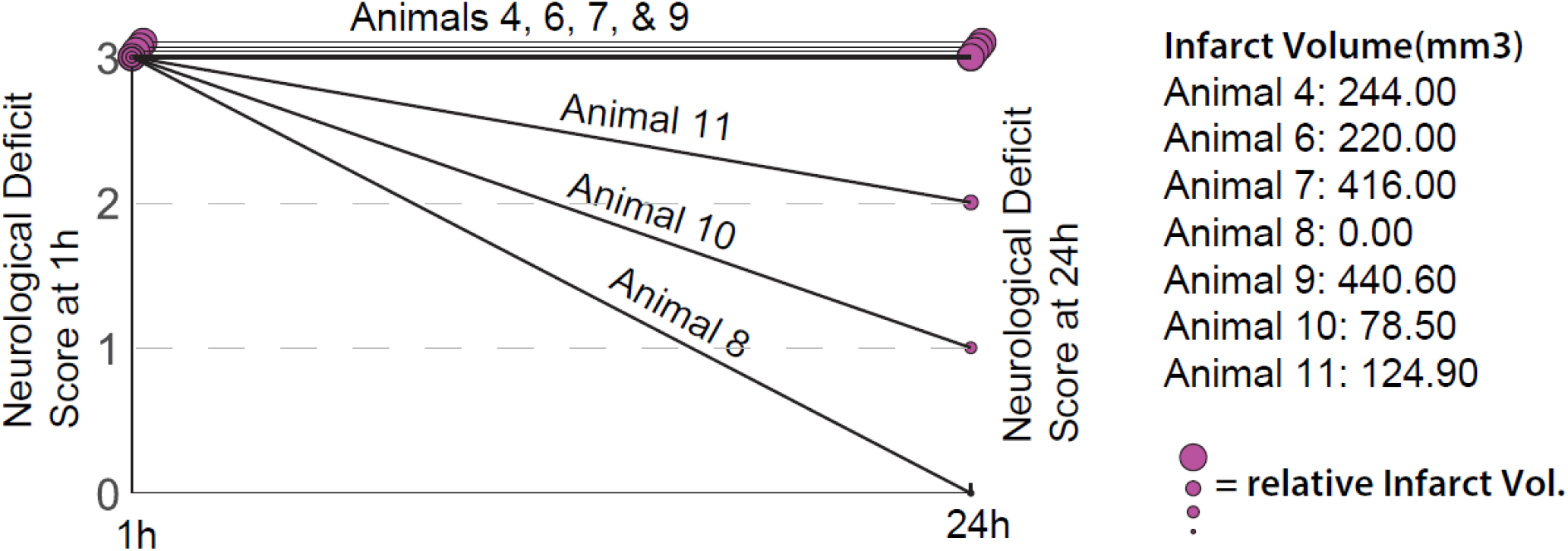
Neurological deficit scores at 24h post-reperfusion was proportional to the infarct volume. The size of the circle (red) represents the relative infarct volume for each animal.

## 4. Discussion

In this study, we used a miniaturized continuous-wave NIRS device to monitor cerebral oxygenation parameters during transient middle cerebral artery occlusion in seven male Sprague Dawley rats. Our optical sensors allowed continuous measurement from both cerebral hemispheres before and during occlusion as well as after reperfusion. Our results indicated that the magnitude of cerebral deoxygenation on the contralateral hemisphere relative to the ipsilateral hemisphere was significantly correlated with the infarct volume.

Previous clinical and animal studies have reported that changes in the contralateral side of cerebral blood supply to the brain are important compensated mechanisms for reducing brain lesion and/or improving neurological function recovery after stroke (Horgan & Finn, 1997). For example, increased blood flow velocity of the contralateral internal cerebral artery and basilar artery in the post-stroke recovery stage was associated with better motor function in rats (Li et al., 2010).

Another study identified that limited extent of native leptomeningeal collaterals affects downstream hemodynamics in Balb/C mice. Lower collateral flow during- and one day after MCAO, coincided with a greater infarct size and worse functional outcome (Kanoke et al., 2020). Our findings further support the notion that adaptive changes of blood supply from the contralateral hemisphere compensates the ipsilateral side after the onset of stroke and leads to smaller infarct formation/or better neurological function recovery. Future study is needed with additional approaches to identify whether native leptomeningeal collaterals or other cerebral artery branches are main components for elevated HbO2 (ΔHbO2).

To our knowledge, this is the first study to use a noninvasive optical imaging tool, namely near infrared spectroscopy (NIRS), to investigate the relationship between cerebral oxygenation and outcome in a preclinical model of ischemic stroke. The NIRS sensors used in this study utilized LEDs and photodiodes, which allowed noninvasive measurements of cerebral oxygenation parameters in rats, unlike other fiber optic-based techniques, which cost significantly more (> 20X) and are invasive and require surgical implantation prior to the experiment.

This proof-of-concept study for using a noninvasive functional imaging tool in a small laboratory animal species, i.e. the Sprague-Dawley rat, provided the first evidence of in vivo assessment of cerebral ischemia using an economical, noninvasive, and easy to apply technique. Continuous-wave NIRS enables longitudinal recording of cerebral oxygenation, which will enhance the rigor of stroke research by enabling better validation of injury and reperfusion. Continuous recording of cerebral oxygenation may also expedite novel fields of stroke research such as spatio-temporal study of pathophysiology of infarct formation and evolution, peri-infarct depolarization (Wolf et al., 1997), real-time CBF monitoring (Culver et al., 2003), and estimation of the hypoxic state of brain cells (Kuo et al., 2014). This technology may also facilitate drug discovery for stroke recovery-enhancing drugs (e.g. monoamine agonists like amphetamines (Stroemer et al., 1998) and neurotrophic growth factors (Kawamata et al., 1997)) that require prolonged assessments.

In conclusion, our results show feasibility of using a noninvasive optical imaging instrument, namely NIRS, in monitoring cerebral ischemia in rodent stroke model. This cost-effective, non-invasive technique will improve the rigor of experimental models of ischemic stroke by enabling in vivo and longitudinal assessment of ischemic injury.

## Acknowledgements

We thank Melanie Roed for technical assistance. Research reported in this publication is based in part upon work supported by 1R43NS100174 and P20GM130443. The content is solely the responsibility of the authors and does not necessarily represent the official views of the National Institutes of Health.

